# Modelling species presence-only data with random forests

**DOI:** 10.1101/2020.11.16.384164

**Authors:** Roozbeh Valavi, Jane Elith, José J. Lahoz-Monfort, Gurutzeta Guillera-Arroita

**Affiliations:** School of BioSciences, University of Melbourne, Parkville, Victoria, Australia

**Keywords:** Class overlap, class imbalance, down-sampling, ecological niche model, Gini index, Hellinger distance, presence-background, recursive partitioning

## Abstract

1. The Random Forest (RF) algorithm is an ensemble of classification or regression trees, and is a widely used and high-performing machine learning technique. It is increasingly used for species distribution modelling (SDM). Many researchers use implementations of RF in the R programming language with default parameters to analyse species presence-only data together with background samples. However, there is good evidence that RF with default parameters does not perform well with such species “presence-background” data. This is often attributed to the typical disparity between the number of presence and background samples also known as *class imbalance*, and several solutions have been proposed.
2. Here, we first set the context: the background sample should be large enough to represent all environments in the region. We then aim to understand the drivers of poor performance of RF with presence-background data, and explain, test and evaluate suggested solutions. Using simulated and real species data, we compare performance of default RF with other weighting and sampling approaches.
3. We show that *class overlap* is an important driver of poor performance, alongside class imbalance. The results demonstrate clear evidence of improvement in the performance of RFs when class imbalance is explicitly managed by sampling methods or when the overfitting commonly associated with overlapping classes is avoided by forcing shallow trees.
4. Presence-background data is a particular version of class imbalance in which class overlap is highly likely and extreme imbalance exists. Without compromising the environmental representativeness of the sampled background, we show several approaches to fitting RF that ameliorate the effects of imbalance and overlap, and allow excellent predictive performance. Understanding the problems of RF in presence-background data allows new insights into how best to fit models, and should guide future efforts to best deal with such data.

## Introduction

Species distribution modelling (SDM) is central to ecological and conservation studies. SDM combines species occurrence data and environmental variables to predict species spatial distributions (Franklin, 2010). Many modelling methods are used for fitting SDMs (Guisan *et al.*, 2017). While earlier examples of SDM mostly focussed on regression methods (Guisan *et al.*, 2002), attention has turned towards the application of machine learning (ML) methods (Elith *et al.*, 2006). This is mainly due to their flexible fitted functions, automatic variable selection methods, ability to handle different types of data and resulting powerful predictive performance (Olden *et al.*, 2008; Hastie *et al.*, 2009). There is a plethora of ML methods available for modelling and mapping species distributions, among which random forests, RF (Breiman, 2001), has become very popular (Cutler et al., 2007; Zhang et al., 2019) because it tends to predict well with little tuning of its parameters (Freeman *et al.*, 2016). RF is also viewed as useful because it can capture complex relationships between species and the environment, it inherently includes interactions in the fitted model (Elith, 2019), and estimates of variable importance are readily available.

In contrast to the often-reported good performance of RF, there are cases where RF has predicted species distributions poorly, often linked to use of presence-background data. Background samples are samples that are randomly taken from the landscape to be used along with presences for modelling species presence-only data. The basic intuitive idea is that this allows SDM methods to compare the environmental conditions favoured by the species to those *available* throughout the landscape under consideration. For instance, Johnson *et al.* (2012) and Shabani *et al.* (2016) compared several SDM modelling methods fitted to presence-background data and found that RF models showed poor predictive performance and overfitting. There are several similar examples in the SDM literature indicating poor performance of RF when fitted on presence-background data. In these studies, the weak performance of RF is attributed to the large number of background samples and authors have suggested approaches for achieving fewer background points (e.g. see Barbet-Massin *et al.*, 2012; Liu *et al.*, 2013). Poor performance has also been reported when modelling presence-absence data containing a large number of absences (Evans & Cushman, 2009; Freeman *et al.*, 2012; Robinson *et al.*, 2018). This behaviour of RF models is well-known within ML, where it is similarly often linked to the sensitivity of these models to imbalanced classes, i.e. where one class (majority class, here background points) substantially outnumbers the other class(es) (minority class, here presence points) (Chen *et al.*, 2004; He & Garcia, 2009; Khalilia *et al.*, 2011).

Although several SDM studies have reported issues with RFs, and have suggested solutions, none focus on explaining the problem and the merits and limitations of the proposed solutions. Here we address this gap, by explaining how RFs work and how common features of SDM data impact their performance. We focus on presence-background data, but also show when to expect problems with presence-absence datasets. We use the ML literature to shed light on the underlying issues. We test how readily available solutions suggested in both the SDM and the machine learning literature work with simulated data, and with a real dataset.

### Random forests: an overview

A Random Forest is an ensemble of classification or regression trees (CART). Classification and regression trees are based on recursive partitioning methods (Box 1). The recursive partitioning repeatedly splits the data into potentially high-dimensional rectangular partitions of the covariate (predictor) space, choosing those for which the response data are relatively homogenous (Strobl *et al.*, 2009). Individual trees are known to have very high variance, i.e. a small change in the training data can considerably change the fitted tree, leading to poor generalisation. An ensemble of many trees, however, generalises better (Hastie *et al.*, 2009). RF fits many trees (also denoted as *base learners* in the ensemble), usually several hundred to thousands, and combines the results of their predictions (Strobl *et al.*, 2009; Elith, 2019). Depending on the response variable, RF uses classification trees (CTs) or regression trees (RTs) for qualitative and quantitative responses, respectively. Each tree is fitted on a bootstrap sample of the training data, which is a random sample, taken with replacement, with total sample size equal to the number of records in the training data. On average, each bootstrap sample contains 63.2% unique records (Efron & Tibshirani, 1994), and these are called in-bag samples. Samples not selected are called out-of-bag samples and are used to estimate the model’s error.

Unlike other closely related ensembling methods such as bagging (Elith, 2019), RF chooses only a random subset of predictors on each split to find the best splitting predictor, instead of searching over all predictors (Box 1). This produces decorrelated trees, so RF tends not to overfit even with many trees (Breiman, 2001; Strobl *et al.*, 2009). The number of predictors selected at each split is represented by the parameter *mtry* in many RF packages. RF by default grows “deep” trees, i.e. unpruned trees with many levels (Strobl *et al.*, 2009; Elith, 2019). This can lead to few observations in terminal nodes, with the absolute minimum allowed varying across packages. In unpruned trees, depth (Box 1) can be controlled by several parameters i.e. minimum number of samples in the terminal nodes, maximum number of terminal nodes or maximum tree depth. The specific options for controlling depth depend on the implementation.

##### Box 1. Tree growing and splitting criteria

Decision trees are grown by splitting the response data on predictor variables. On a chosen predictor each split point is selected to separate the response into two nodes, aiming to achieve the highest purity/similarity within each node, and the highest impurity/dissimilarity between nodes. On each split, all predictors are tested and the predictor causing the highest impurity gain is chosen. Each level of splits is a tree depth. The impurity is calculated with measures such as the *Gini index* for classification trees (CTs). For binary data and CTs, the Gini index is calculated as 2*p*(1 − *p*), where p is the proportion of one of the classes (e.g. in species data, presence and absence/background are the classes) (Hastie et al., 2009). The splitting criterion for regression trees (RTs) is usually calculated using a sum of squares instead of the Gini index. Taking the average of the response y as the predicted value 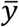 in each node, sum of squares is calculated as 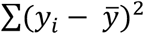 over all *i* observations (Hastie *et al.*, 2009).

Breiman (Breiman, 2001, 2002) proposed the Gini index as the default splitting criterion for CTs (as single trees and within RFs). The Gini splitting criterion is now known to have weaknesses when applied to datasets with high class imbalance (Drummond & Holte, 2000; Cieslak & Chawla, 2008). Alternative criteria have been developed that are relatively insensitive to class imbalance such as *Hellinger distance* (Cieslak & Chawla, 2008). The Hellinger distance was first implemented in decision trees by Cieslak and Chawla (2008), known as Hellinger distance decision tree (HDDT). This method has been shown to be a robust criterion and insensitive to class imbalance and is claimed to be superior to the traditional splitting criteria such as Gini when used on imbalanced data (Cieslak *et al.*, 2012).

#### Prediction with random forests for binary data

Here we focus on how the predicted values are estimated in RFs, since binary data can be treated in two ways. If binary data are treated as classes (hereafter named classRF), for any one tree the algorithm predicts the outcome class in a terminal node as the most frequent class across the observations in that node (also known as *majority vote*). Each such predicted outcome of a terminal node is called a vote (Breiman, 2001; Strobl *et al.*, 2009). The classRF prediction class for any observation is then calculated again by majority voting, now across all fitted trees. For any prediction, class probabilities can also be calculated (Liaw & Wiener, 2002) as the proportion of the predicted positive (“1”) class of all CTs in the final classRF model (Breiman, 2002). Binary data can also be modelled with RTs (Malley *et al.*, 2012; Zhang *et al.*, 2019), hereafter called regRF. This treats the response as continuous, and the mean over all observations in a terminal node is the predicted node outcome, which is then averaged over all trees in the final regRF model (Hastie *et al.*, 2009). In regRFs, the terminal node mean is equivalent to proportion of presences (1s) in that node (Zhang *et al.*, 2019). Given the different splitting criteria and calculation of predictions across trees, predicted probabilities for binary responses will not necessarily be identical between RTs and CTs (Malley *et al.*, 2012). Both prediction approaches are implemented in the R package *randomForest* (Liaw & Wiener, 2002). Whilst most modellers use classRF for binary data, Malley *et al.* (2012) found that probabilities estimated by regRF are more accurate and consistent than those from classRF (Malley *et al.*, 2012).

The R package *ranger* (Wright & Ziegler, 2017) is a fast implementation of RFs. For modelling binary data, they use the ideas of Malley *et al.* 2012, naming the resulting regRF as a *probability forest*. Instead of using the traditional sum of squares split criteria, for probability forests they use the Gini index or Hellinger distance for splitting. As described for regRF above, predictions are estimated as class proportions at terminal nodes (response mean) then averaged across all trees (Wright & Ziegler, 2017).

### Understanding class imbalance

The predictive performance of RF and many other ML classifiers can be substantially affected by class imbalance. There has been considerable interest in reasons and solutions for these impacts in the ML literature, for both RF and other algorithms (Chen *et al.*, 2004; He & Garcia, 2009; Ali *et al.*, 2015; Krawczyk, 2016). The obvious issue with class imbalance is unequal representation of the classes, and this is known to cause problems (He & Garcia, 2009; Khalilia *et al.*, 2011). However, other issues also occur with imbalanced data, including small sample size of the minority class, small disjuncts within the minority class, and class overlap (Japkowicz, 2003; Denil & Trappenberg, 2010; Ali *et al.*, 2015). We do not expand on the small samples and small disjuncts here because we expect they are less relevant to the SDM presence-background problem than imbalance and overlap.

Class overlap occurs in covariate space, when two classes co-occur over the same multi-covariate ranges. It is also referred to as class complexity or class separability (Japkowicz & Stephen, 2002; Prati *et al.*, 2004). Class overlap is often the most serious problem in imbalanced data (Prati *et al.*, 2004; Denil & Trappenberg, 2010; Ali *et al.*, 2015). When classes overlap in some or even all of the covariate space, it is very hard to determine discriminative rules to separate the classes (Ali *et al.*, 2015). The splitting procedure of RF tries to separate classes that may not be completely separable with the available predictors. Since a RF model by default produces very deep trees, the algorithm continues to attempt to separate the classes to reduce impurity and may go to the level of individual observation, thus overfitting to the training data.

#### A demonstration with presence-absence data

Class overlap will by definition occur in species presence-background data, because the background points are meant to sample the full environmental space in the region, including that in which the species occurs (Renner *et al.*, 2015). However, it can also occur in presence-absence data if both presences and absences are recorded in similar environmental conditions. This can occur, for instance, in species that are sparse (low abundance) over a large range of their prime habitat (Rabinowitz, 1981). Since presence-absence data are simpler to conceptualise, we start here with simulated data showing impacts of class overlap in RF models.

We simulated two presence-absence datasets for two virtual species (Figure 1, and code in Appendix S3). Both these datasets are severely imbalanced (species prevalence below 0.06, i.e. less than 6% of presence class) and both species occur in only a small range of the predictor variable. Species 1 (Figure 1a) has low occurrence probability in “suitable environments” (defined as the best environmental conditions), so the species could be absent in many parts of its habitat. Hence there will be both presences and absences within the same “suitable” environmental condition leading to extensive class overlap. In contrast, species 2 (Figure 1c) has high prevalence within the narrower range of suitable environments, leading to minimal class overlap (very few absences recorded within “suitable” environments). We generated 50 datasets of 300 records (presence and absence) from each species. We model these using a RF through the randomForest package with default settings (classRF, fitting 500 trees to all records) and also with an implementation of RF that handles class imbalance by down-sampling the majority class (explained in following sections). Predictive performance is estimated on 1500 independent records for each species.

**Figure 1:**
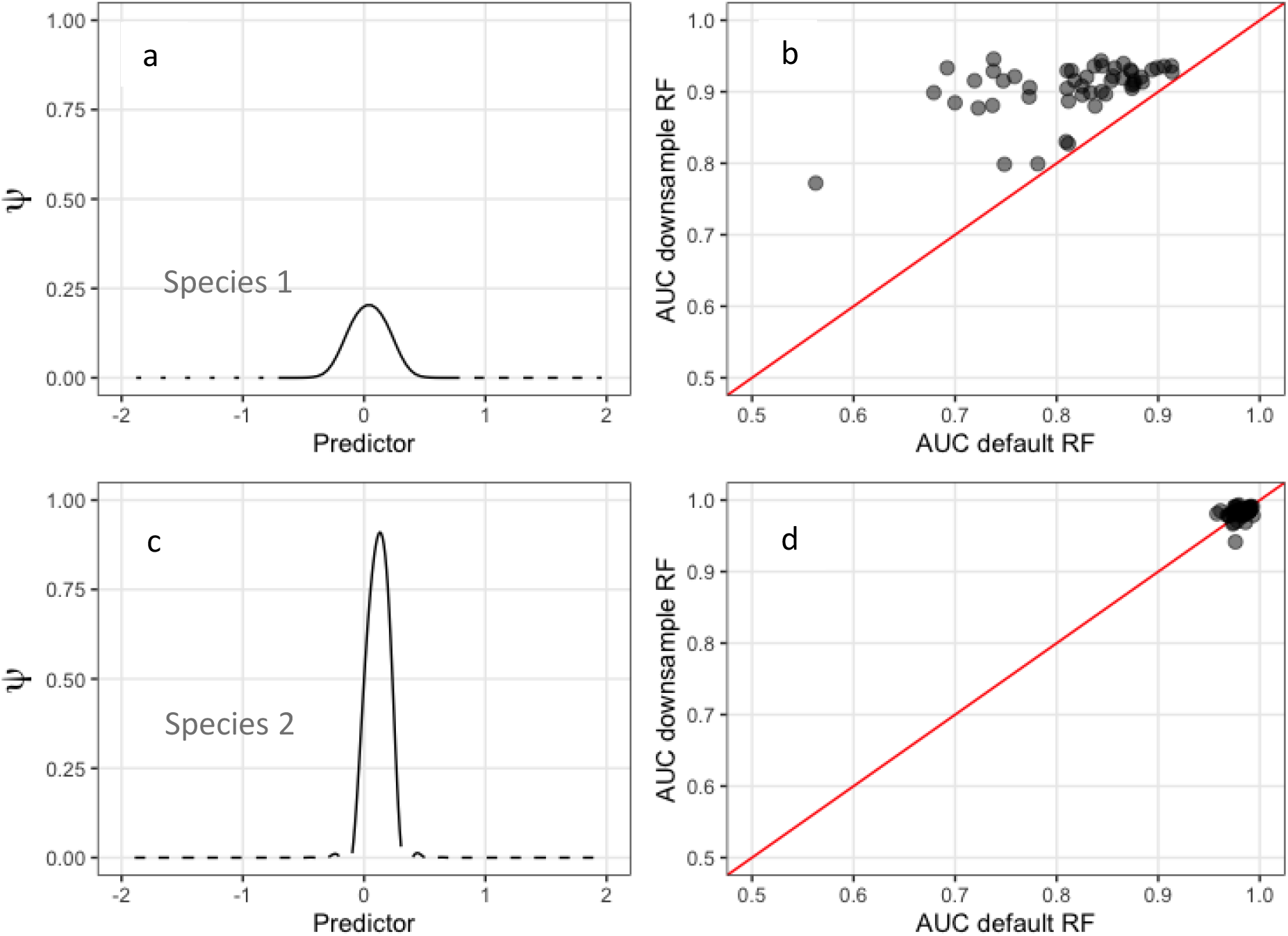
Two species distributions simulated for assessing the impact of class overlap on RF. The left plots (a & c) show the probability of species occurrence (ψ) as a function of a single predictor. We assume the predictor is equally represented across the landscape. Plots b & d show the predictive performance of RF with default settings vs RF using a method for dealing with class imbalance (down-sampling, see section “suggested solutions”). See “data availability” section for code.

For species 1, neither approach to fitting RFs achieves an AUC very close to one (as expected for such distribution characteristics), but the performance of the down-sampling approach (AUC ranging from 0.78 to 0.94) is always better than that of the default RF (AUC from 0.57 to 0.92, Figure 1b). This demonstrates the gain achieved from improving model fit by effectively equalising class sizes (see next section). In contrast, for species 2 with high probability of presence within its preferred environmental conditions (Figure2, c; little class overlap yet still very low prevalence across all predictor values and hence high class imbalance), both approaches to fitting a RF model give AUCs close to 1 (Figure 1, d). The contrast between results for species 1 and species 2 shows that it is not the imbalance of classes itself that impacts predictive performance, but class overlap. This simple example illustrates that when classes can be clearly separated by the available environmental conditions, commonly used default methods for fitting RF models perform well, regardless of class imbalance. Similar results are reported by Japkowicz and Stephen (2002) that decision trees (and two other ML algorithms) are less sensitive to class imbalance when classes are separable.

**Figure 2.**
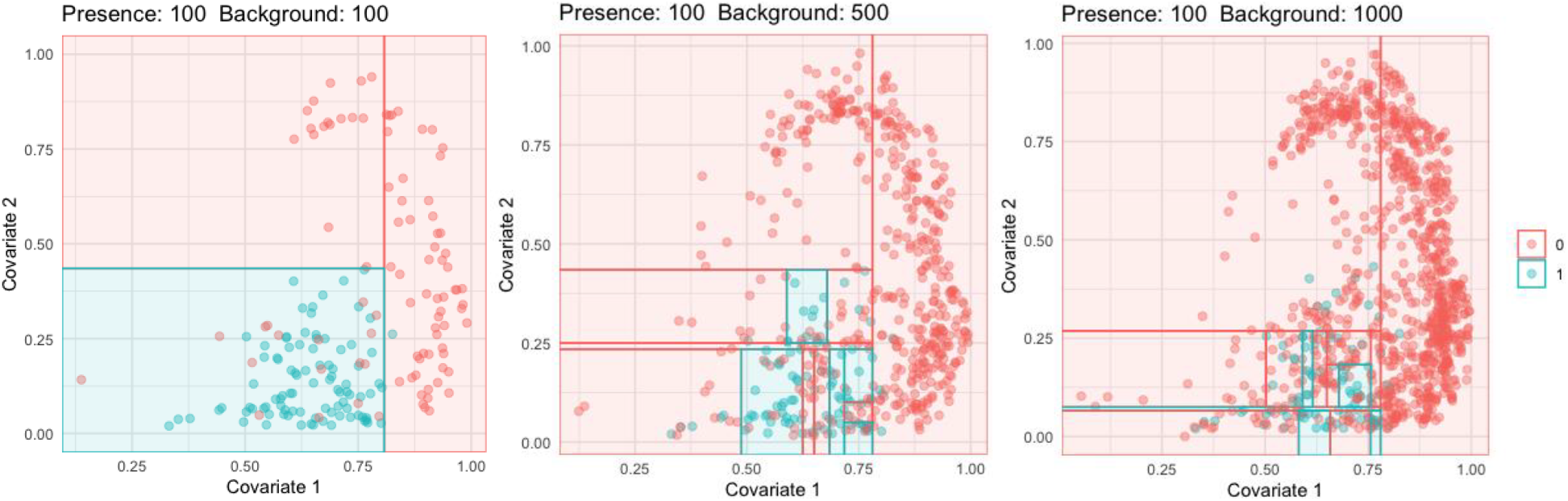
Visualisation of recursive partitioning with different number of background points. The 0s (red) and 1s (blue) show background and presence samples, respectively. The blue sections are the terminal nodes predicted as presence and red are predicted as background. The lines in the plots show the split points. This is a tree fitted by *rpart* package (Therneau *et al.*, 2019) with maximum allowed depth of 25 and complexity parameter of 0, i.e. no early stopping imposed.

#### The problem with presence-background data and RF

In presence-background data the two classes are presence and background, and class overlap is expected because the background samples are meant to sample all environments, including those preferred by a species. With background samples obtained randomly and independently of species locations, there is potential for exact spatial and therefore environmental overlap (i.e. background points can be at the same location as presence points). Furthermore, background samples are meant to characterize environmental conditions in the region of interest, so potentially tens of thousands of samples are needed, depending on the complexity of environments in the region (Franklin, 2010; Renner *et al.*, 2015), leading to severely imbalanced datasets. So, when modelling presence-background data, it is common to end up in the dreaded combination of sever class imbalance and class overlap. The solution is not to sample fewer background points (risking not properly representing the background environmental conditions) or to ensure that presence and background samples do not co-occur, because all that does is to move away from the statistically correct way to create these samples. A different solution is needed.

To understand how tree-based models fit presence-background samples, we simulate presence and random background samples across 2 covariates (Figure 2; see “data availability” section for code). We fit CTs to the data, with the algorithm allowed to generate a deep tree if warranted by the data (similar to RF that fits deep trees). With a 1:1 ratio of presence to background (left panel), despite some class overlap, the CT stops at a simple two-split model because no improvement can be made by further splits. As more background points are added, more overlap and more imbalance is inevitable, so the algorithm continues splitting out smaller and smaller areas where presences are still dominant. Note here that, with presence-background data, class imbalance and class overlap both necessarily come into play. It has been shown, using other ML algorithms, that imbalance and overlap tend to interact, and result in overly complex models (Denil & Trappenberg, 2010). That is likely occurring here. Since we know from theory that many background samples are needed to properly represent the background environmental conditions, what is the solution?

### Suggested solutions

Here we focus on solutions relevant to RF fitted to presence-background data. We include approaches suggested by both SDM and ML researchers. Whilst there are several proposed solutions in the ML literature, we only explore methods readily available in R packages, and also those that suit the problem of modelling spatially explicit presence-background data (see discussion). Most solutions focus on tweaking the model or resampling the data.

#### Applying weights

This approach deals with the issue of class imbalance and arose in the ML literature. The idea behind weighting different classes comes from the notion of cost-sensitive learning, and typically works by increasing the misclassification cost on the minority class (Chen et al., 2004). A higher weight (i.e. higher misclassification cost) will be given to the minority class. Weights are estimated in different ways across software implementation of RFs. We use the popular *randomForest* package (Liaw & Wiener, 2002), in which the weighting is applied to the Gini index, impacting selection of splits and also weighting across terminal nodes in each tree (Chen *et al.*, 2004). The weights (cost) are applied to the Gini index (denoted as G) as follows (Breiman *et al.*, 1984; Kuhn & Johnson, 2013):

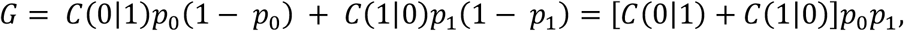

where *p*_0_ and *p*_1_ are the proportion of background and presence classes in the partition respectively, *C*(0|1) is the cost of mistakenly predicting a class 1 (presence) sample as class 0 (background), and *C*(1|0) is the opposite. In the randomForest package the class weights are only applicable for classRF since regRF does not use the Gini index nor does it view the data as classes. The weights we used are inversely proportional to the class distribution in the complete dataset i.e. 1 / *proportion*(*class*_*i*_) (Albon, 2018).

#### Equal-sampling

This idea arose in the SDM literature, in the context of presence-background modelling. The idea of equal-sampling (Barbet-Massin *et al.*, 2012) is to fit ten different RF models on ten different data sets, each with all presences and a random sample of background of size equal to the number of presences. The final prediction is an average of those from the 10 models. This is based on “repeated random sub-sampling”, a technique from the ML literature for dealing with imbalanced classes (Japkowicz & Stephen, 2002; Khalilia *et al.*, 2011). It works by reducing the number of background samples (the majority class) and thus reducing the opportunity for class overlap. Since the number of background samples is matched to the number of presence samples in each repeat, the total number of background samples used in the final model will be 10 times the number of presences. For small presence samples, this has the undesirable consequence that relatively few background samples are taken.

#### Down-sampling

Down-sampling is similar to the equal-sampling approach, but it differs in that it subsamples at the level of the individual tree. It was first suggested in the ML literature (Chen *et al.*, 2004) then used for SDM (Evans & Cushman, 2009; Freeman *et al.*, 2012). It uses the same number of background samples as presence points in each CT in a classRF model. Since forests typically have hundreds of trees, overall this approach uses many more background samples than the equal-sampling technique (above) as each individual tree uses potentially a different set of background samples. Down-sampling works by balancing the training data (Chen *et al.*, 2004) and thus reducing the chance of class overlap and imbalance for each individual tree. Down-sampling is often referred to as balanced RF (Chen *et al.*, 2004; Robinson *et al.*, 2018). This version of RF has been used in several ecological studies to address the challenge of class imbalance in the analysis of species presence-absence data (Evans & Cushman, 2009; Freeman *et al.*, 2012; Robinson *et al.*, 2018). Additionally, in a comparison study on 225 species presence-background datasets, we found that down-sampling RF was among the best performing models (whilst the default RF was the worst model) (unpublished data).

#### Using regRF instead of classRF

The approaches described above use classRF reflecting the fact that many ML techniques model binary data as a classification task, so attention has been directed to solutions for classification methods (Krawczyk, 2016). However regression has also been discussed for binary data (e.g. Malley *et al.*, 2012). As explained in the section on prediction from RFs, Malley *et al.* (2012) provide evidence that class probabilities are better predicted via regRFs rather than classRFs. Likewise, Zhang *et al.* (2019) recommend using regRF with presence-background data. Here regRF is implemented using the randomForest package with presence-background data as a numeric 1-0 response. No weights are used on samples, and all presence and background points are used.

#### Using shallow probability trees (with Hellinger distance splitting)

Our visualisation in Figure 2 shows that trees will tend to try to separate overlapping classes, and with large background samples this implies trees will become deeper and deeper – i.e. the model will become increasingly complex. Here we propose an approach that deals directly with this issue, focussing on the issue of class overlap. Our idea is to enforce shallower trees. In addition, we suggest using a splitting criterion known to be robust to imbalanced data. Within the options of commonly available R packages, this suggests using Hellinger distances in preference to the Gini index (Drummond & Holte, 2000; Cieslak & Chawla, 2008; Cieslak *et al.*, 2012). One way to bring these ideas together is to fit a RF in the *ranger* package (Wright & Ziegler, 2017), since this enables probability forests and control of tree depth. In this approach, no subsampling is required, and all background samples are used in each probability tree of the forest.

## Tests with real and simulated data

## Materials and methods

### Species data for modelling

To assess the predictive performance of these approaches, we used a global dataset (hereafter “the NCEAS dataset”) of 225 species from six regions of the world, fully described and made available by the NCEAS data group (Elith *et al.*, 2020). This dataset has presence-only data for model fitting and presence-absence data from designed surveys for model evaluation. It comprises birds and plants of the Australian Wet Tropics (AWT); birds of Ontario, Canada (CAN); birds, plants, mammals and reptiles of north-east New South Wales, Australia (NSW); plants of New Zealand (NZ); plants from five countries of South America (SA); and plants of Switzerland (SWI). The number of presence-only records ranges from 5 to 5822 per species. From 7 to 12 environmental predictors were used for model fitting. The evaluation datasets range from 102 (AWT) to 19120 (NZ) survey sites (Elith *et al.*, 2020). Since the presence-only data are known to be biased (Phillips *et al.*, 2009), we aim to deal with that bias in our modelling. Rather than using a random background sample, we aim to create a background sample with similar biases to that in the presence-only records. Following Kujala *et al.* (2015), we generated 50,000 background samples from a bias layer constructed using kernel density estimation on the target-group samples (i.e. all species in each biological group) defined by Phillips *et al.* (2009) (see full description and code in Appendix S1).

In addition, we generated 100 virtual species with the *virtualspecies* package v1.5.1 (Leroy *et al.*, 2016). Real environmental variables from Australia were used to simulate these species. We generated a species suitability map by applying a gaussian function (with random mean and standard deviation for each species) on the first four principal component axes of 21 environmental covariates (see “data availability” section for code). The final suitability was a multiplicative function of responses to the four predictors. Then rasters of presence-absence species data for modelling and evaluation were randomly drawn following a binomial distribution with suitability values at each pixel as success rates. Depending on the prevalence of each simulated species, from 20 to 182 presence points were randomly drawn as presence-only modelling data. In addition, 50,000 random background samples of the landscape were taken, and used for all species. For evaluating predictions, 5000 random presence-absence samples were drawn for each species. These were specified to exclude all training set presence sites and their direct neighbours (cells).

### RF implementations

We implemented all the strategies described above (‘Suggested solutions’) alongside a *randomForest* classRF with default parameters. The models were all implemented in the R programming language (R Core Team, 2019) using two R packages, *randomForest* v4.6-14 (Liaw & Wiener, 2002) and *ranger* v0.12.1 (Wright & Ziegler, 2017). The parameters and implementations are described in Table 1, and code is provided in Appendix S1.

**Table 1:** Implemented RF model parameters and descriptions. The training data for all models were the same except for the equal-sample model, which was run 10 times with a reduced background sample.

| RF Model | Parameters | Value | Description | R package |
| --- | --- | --- | --- | --- |
| Default (classRF) | ntree | 1000 | number of trees | randomForest |
| regRF | ntree | 1000 | number of trees | randomForest |
| Weighted (classRF) | ntree | 1000 | number of trees | randomForest |
| | classwt | inverse weights | class weights: inverse weights were calculated as $1 / proportion(class_i)$. | |
| Equal-sample (classRF) | ntree | 1000 | number of trees<br>model is run 10 times with equal number of presence and background samples and predictions are averaged | randomForest |
| Down-sample (classRF) | ntree | 1000 | number of trees | randomForest |
|  | sampsize | as the number of presences | size of the sample to draw from each class |  |
|  | replace | TRUE | take bootstrap sample with replacement |  |
| Shallow (regRF or probability forest) | num.trees | 2000 | number of trees | ranger |
|  | splitrule | hellinger | Hellinger distance splitting criterion |  |
|  | max.depth | 2 | maximal tree depth (=maximum 4 terminal nodes) |  |
|  | probability | TRUE | use ranger probability forest |  |
|  | replace | TRUE | take bootstrap sample with replacement |  |
| Shallow-tuned (regRF or probability forest) | num.trees | 2000 | number of trees | ranger |
|  | splitrule | hellinger | hellinger distance splitting criterion |  |
|  | max.depth | 1 to 4 | maximal tree depth |  |
|  | probability | TRUE | use ranger probability forest |  |
|  | replace | TRUE | take bootstrap sample with replacement |  |

We did not tune all the parameters of the models as RFs are known to work reasonably well with default parameters (Freeman *et al.*, 2016; Probst *et al.*, 2019). For the shallow probability trees, we used the package ranger and chose “maximum depth” of 2 (i.e. maximum 4 terminal nodes; column 1, Table 1). Since the choice of 2 is subjective, to assess the sensitivity of this parameter, we fitted another shallow model and tuned the maximum depth (allowing depths of 1 to 4) using a 5-fold cross-validation on the presence-background training data. As the aim to keep the trees shallow is to avoid overfitting, we did not try values higher than 4 (2^4^ = 16 possible terminal nodes). In comparison, the maximum depth for trees in the randomForest implementations can be very high. This can lead to only 1 observation per terminal node in classRF and 5 for regRF.

### Model evaluation and comparison

Two threshold-independent evaluation metrics were used to assess the predictive performance of the models: Area under the ROC curve (AUC) and Pearson correlation (COR). AUC is a very well-known metric in SDM. It reflects the discrimination power of the models i.e. the ability of the model to discriminate presences from absences (Pearce & Ferrier, 2000). COR values were calculated 1) between the predicted relative likelihood of occurrence and the presence-absence records of the evaluation data, in the NCEAS dataset; and 2) between the predicted likelihood of occurrence and the simulated probability of occurrence, for the virtual species. Note that we are not including metrics that directly measure calibration, because presence-background methods cannot be properly calibrated by nature of the data inputs (Guillera-Arroita *et al.*, 2015).

To determine whether the modelling approaches had statistically different performance, we used non-parametric hypothesis tests suggested for comparison of models over multiple datasets (Demšar, 2006; García & Herrera, 2008). The Friedman’s Aligned Rank test (García & Herrera, 2008) was calculated for AUC (and COR) and the p-values were adjusted by Bergmann-Hommel correction (Bergmann & Hommel, 1988). All pairwise comparisons were assessed to check whether the differences between different approaches and default models were statistically significant.

## Results

### Performance on real dataset

The performance of the models on the real dataset (the NCEAS data) is shown in Figure 3. Using 0.7 as the cut-off for useful discrimination (Pearce & Ferrier, 2000), the default RF had poor performance in all regions except SWI. It was the poorest performer overall, followed by weighted and regRF. The down-sample, equal-sample and both shallow RF models performed noticeably better for both AUC and COR, for all regions (Figure 3). The Friedman’s Aligned Rank test was highly significant, indicating the statistical difference between the models in both metrics. The adjusted p-values showed: 1) the difference in down and equal samples is not significant, and they tend to be slightly better than both shallow models 2) shallow and shallow-tuned are not significantly different, 3) all these four models are significantly better than the default, weighted and regRF (see Appendix S1; Figures S4, S5 and S6). The values of tree depth chosen by the tuning favoured depth of 4 (190 species were fitted with depth 4 and only 8 with depth 2).

**Figure 3.**
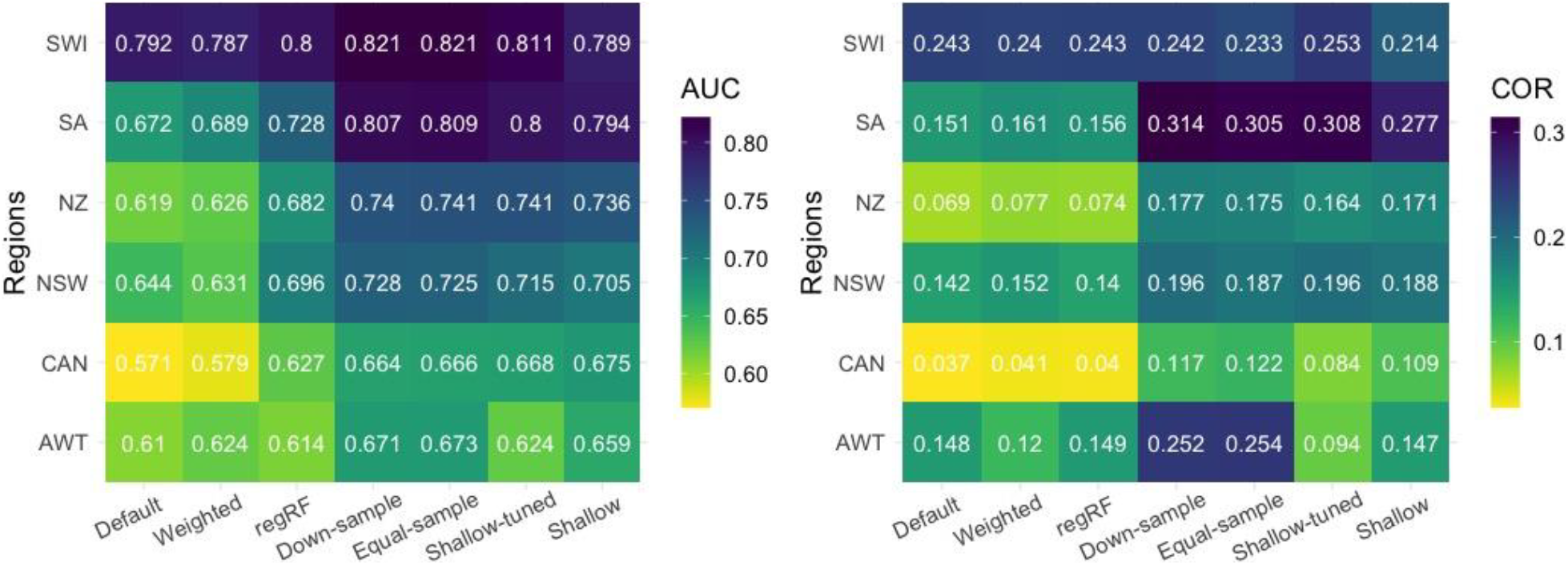
Performance of different RF models on different regions of NCEAS data. The values in cells are AUC and COR for the left and right plots respectively. Colour varies with values, yellow (low) to dark navy (high).

These trends in performance were generally consistent across regions, though the differences between methods in some regions (e.g. SA) were starker than in others (e.g. SWI, where differences were minor and did not demonstrate the general trends, e.g. for COR).

### Performance on simulated dataset

Figure 4 shows the performance of the models on 100 simulated species as evaluated with AUC and COR. The default model had poor performance on both evaluation metrics, with a median AUC ~0.62 and median COR less than 0.2. The weighted classRF had a very similar performance and showed no improvement. The two (equal and down) sampling approaches and the shallow RFs clearly improved performance (Figure 4), achieving median AUC higher than 0.85 and median COR higher than 0.75. Both shallow approaches produced slightly better results than other models. Across all species the selected tree depth in the shallow-tuned model varied from 1 to 4, with similar numbers of each (25, 21, 25 and 29 times for depth 1-4, respectively). However, the tuned version of shallow RF showed no evident performance boost over the shallow with constant max-depth of 2. The regRF showed much better discrimination than default and weighted classRF, but very similar COR. This is in line with the result found by Zhang *et al.* (2019). The spatial prediction of implemented methods on an example virtual species is presented in Figure S10 of Appendix S1.

**Figure 4:**
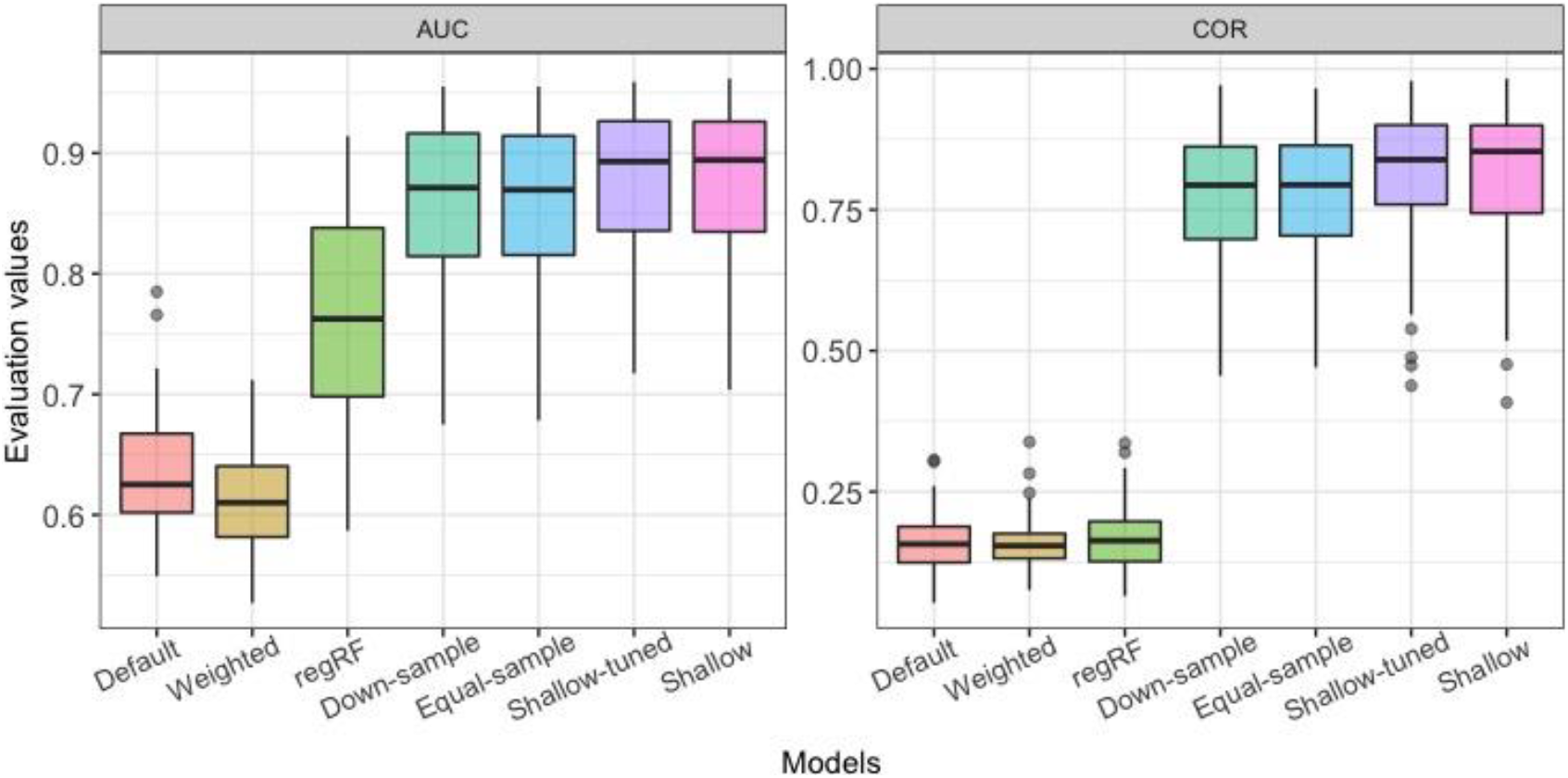
Performance of different RF models on 100 virtual species. The left plot is the AUC and right one is COR values.

The Friedman’s Aligned Rank test was highly significant, indicating the statistical difference between the models. The p-values in AUC showed: 1) shallow and shallow-tuned are not significantly different, 2) they are significantly better than down-sample and equal-sample. These four models are not significantly different from one another in COR. Down-sample, equal-sample and both shallow RFs are significantly better than other methods in both metrics (Appendix S1).

## Discussion

We have explained the problems facing RF models fitted to presence-background data and explored the predictive performance of suggested solutions proposed in the ML and SDM literature. Several solutions (equal-sampling, down-sampling and shallow RF) improved performance significantly. Here we focus on those solutions, interpreting their approach and performance in the light of the presence-background modelling problem.

### Effectiveness of sampling methods for class imbalance

Down-sampling and equal-sampling provide similar results, and both substantially improved predictions in both real and simulated datasets. Both deals directly with the amount of class imbalance and overlap by sub-sampling the majority class (background), but they do it differently. Down-sampling focusses on data used in each tree, in contrast to equal-sampling which provides sub-sampled sets at the level of the forest, repeated. This means that down-sampling is faster when it comes to prediction to rasters and covers more of the full set of background samples. Based on theory, methods that use more background samples are preferable, up to the point that all environmental space in the region is well represented. Indeed, for all presence-background models there are good statistical reasons for providing enough background samples to cover the environmental space (usually tens of thousands) and to sample them irrespective of presence locations (Warton & Shepherd, 2010; Renner *et al.*, 2015). This implies that – even though there is no noticeable performance benefit here of down-sampling over equal-sampling, one would expect that in some situations the additional environmental information in larger samples would be beneficial. Given its speed and easy implementation, we believe down-sampling is a good choice.

Resampling is commonly used for handling imbalanced data in the ML literature for methods such as RF and decision trees (Cieslak *et al.*, 2012), but SDM users have not adopted it as widely. Where used, several examples show benefits of down-sampling (Evans & Cushman, 2009; Barbet-Massin *et al.*, 2012; Robinson *et al.*, 2018; Shaeri Karimi *et al.*, 2019). However, improvement is not universal. For instance, Freeman *et al.* (2012) evaluated the effectiveness of down-sampling on presence-absence data of tree species distributions. They did not find any improvement with the down-sampling technique over default RF. In this case there may have been high separability between their species presence and absence data (i.e. low class overlap).

Other methods of resampling have been proposed in the ML literature, taking the opposite approach and up-sampling the minority class (Kuhn & Johnson, 2013). Since with presence-background data the true data are the presences and the background samples are simply a technique for fitting the models, we chose to leave the true data untouched and only resample background points.

### Another approach of handling class imbalance in RFs

In this study we also tested a new approach, based on our understanding of the dual impacts of class imbalance and class overlap, and our desire to use all background samples. In this approach, we used probability forest (regRF) with shallow trees and Hellinger distance splitting criterion (the shallow models) without any subsampling i.e. all background samples were used in model fitting. The shallow method had a substantial improvement over the default classRF model. We avoided fitting deep trees by limiting the maximum depth of the trees in this model. The result showed tuning did not significantly improved over the subjective choice of 2 for max-depth parameter (table 1) in shallow model. Hence, this approach shows promise but needs further investigation for performance across datasets because it moves away from the usual approach of using deep trees in RFs.

### Final remarks

When applied on presence-background species distribution data, the default implementation of RF model shows relatively poor predictive performance. RF and in general decision trees focus on separating different classes (or similar continuous data), so they have an intrinsic issue with data that have overlap between classes, such as species presence-background data. This problem happens in RFs with both classification and regression trees. We show RF can be used effectively to fit SDMs by adjusting the implementation to accommodate the realities of the data (different proportion of classes that overlap). There are several approaches that effectively handle the problem.

## Acknowledgments

Roozbeh Valavi was supported by an Australian Government Research Training Program Scholarship and a Rowden White Scholarship.

## Authors’ contributions

Roozbeh Valavi conceived the initial idea, ran the analyses, and led the writing of the manuscript. All authors contributed critically to the discussion of ideas and manuscript drafts.

## Data Availability

All environmental raster data and simulation codes will be available on Open Science Framework (OSF).

